# TeXP: Deconvolving the effects of pervasive and autonomous transcription of transposable elements

**DOI:** 10.1101/648667

**Authors:** Fabio CP Navarro, Jacob Hoops, Lauren Bellfy, Eliza Cerveira, Qihui Zhu, Chengsheng Zhang, Charles Lee, Mark B. Gerstein

## Abstract

Long interspersed nuclear element 1 (LINE-1) is a primary source of genetic variation in humans and other mammals. Despite its importance, LINE-1 activity remains difficult to study because of its highly repetitive nature. Here, we developed and validated a method called TeXP to gauge LINE-1 activity accurately. TeXP builds mappability signatures from LINE-1 subfamilies to deconvolve the effect of pervasive transcription from autonomous LINE-1 activity. In particular, it apportions the multiple reads aligned to the many LINE-1 instances in the genome into these two categories. Using our method, we evaluated well-established cell lines, cell-line compartments and healthy tissues and found that the vast majority (91.7%) of transcriptome reads overlapping LINE-1 derive from pervasive transcription. We validated TeXP by independently estimating the levels of LINE-1 autonomous transcription using ddPCR, finding high concordance. Next, we applied our method to comprehensively measure LINE-1 activity across healthy somatic cells, while backing out the effect of pervasive transcription. Unexpectedly, we found that LINE-1 activity is present in many normal somatic cells. This finding contrasts with earlier studies showing that LINE-1 has limited activity in healthy somatic tissues, except for neuroprogenitor cells. Interestingly, we found that the amount of LINE-1 activity was associated with the with the amount of cell turnover, with tissues with low cell turnover rates (e.g. the adult central nervous system) showing lower LINE-1 activity. Altogether, our results show how accounting for pervasive transcription is critical to accurately quantify the activity of highly repetitive regions of the human genome.

**Author Summary:** Repetitive sequences, such as LINEs, comprise more than half of the human genome. Due to their repetitive nature, LINEs are hard to grasp. In particular, we find that pervasive transcription is a major confounding factor in transcriptome data. We observe that, on average, more than 90% of LINE signal derives from pervasive transcription. To investigate this issue, we developed and validated a new method called TeXP. TeXP accounts and removes the effects of pervasive transcription when quantifying LINE activity. Our method uses the broad distribution of LINEs to estimate the effects of pervasive transcription. Using TeXP, we processed thousands of transcriptome datasets to uniformly, and unbiasedly measure LINE-1 activity across healthy somatic cells. By removing the pervasive transcription component, we find that (1) LINE-1 is broadly expressed in healthy somatic tissues; (2) Adult brain show small levels of LINE transcription and; (3) LINE-1 transcription level is correlated with tissue cell turnover. Our method thus offers insights into how repetitive sequences and influenced by pervasive transcription. Moreover, we uncover the activity of LINE-1 in somatic tissues at an unmatched scale.

## Introduction

Long interspersed nuclear element 1 (LINE-1) has attracted much attention in the last decade due to its capacity to promote genetic plasticity of the human genome. LINE-1 is a DNA sequence capable of duplicating itself and other DNA sequences by mobilizing messenger RNAs (mRNAs) to new genomic locations via retrotransposition [1–3]. There are multiple molecular mechanisms to deactivate LINE-1 instances, most prominently, the truncation of 5’UTR due to partial retrotransposition has resulted in mostly inactive and truncated copies of LINE-1 across the human genome[3–6]. Truncated copies of LINE-1 lack their internal promoter sequence and therefore, are expected to be dead-on-arrival. Although full-length LINE-1 activity has been described in both healthy and pathogenic tissues [3,7,8], quantifying its activity is remarkably difficult due to its repetitive nature. Until recently, LINE-1 retrotransposition was believed to occur in germ cells [9–11] and tumors [12–14], but not in somatic tissues. However, growing evidence suggests that LINE-1 is active in the neuroprogenitor cells and in other healthy somatic tissue at low levels [15–18].

As opposed to healthy tissues, tumor and tumor derived cell lines show higher levels of LINE-1 activity [13]. LINE-1 instances are likely to be activated due to broad demethylation of LINE-1 promoter [19]. The literature describes many other factors contributing to the constraints of LINE-1 activity pre- and post-transcriptionally [20]; however, little is known about its activation and impact in tumors [21]. A major challenge to asses LINE-1 activity is the requirement of either specialized assays [22,23] or multiple and complementary datasets [24], hindering estimation of autonomous LINE-1 transcription in a large number of samples. Moreover, affordable methods to quantify LINE-1 activity, such as those based on RNA [17,25,26], are largely confounded by the high copy number nature of LINE-1 and pervasive transcription [23], which refers to the idea that the majority of the genome is transcribed, beyond just the boundaries of known genes [27].

How much pervasive transcription influences the human transcriptome is still unclear [27–29]. Some researchers suggest that pervasive transcription is mostly derived from technical and biological noise and, therefore, might not be relevant in RNA sequencing experiments [30]. Others suggest that pervasive transcription has a stochastic nature, and if sequenced at enough depth the majority of the genome may be transcribed. With either theory, pervasive transcription should not affect quantification of the transcription of protein coding genes, which are present either in single copy or low copy numbers in the genome. However, the quantification of the transcriptional activity of transposable elements, including LINE-1, would be greatly affected by pervasive transcription due to their multi-copy nature. The autonomous transcription of LINE-1, on the other hand, derives from LINE-1 transcripts being fully transcribed from its internal promoter. Thus, by definition, since LINE-1 promoters are at the 5’ extremity of LINE-1 elements, autonomous transcription is more likely to derive from full length LINE-1 instances. These transcripts could derive from both intronic or intergenic full-length LINE-1 instances.

This paper presents a new method to remove the effect of pervasive transcription on RNA sequencing datasets and reliably quantify LINE-1 subfamily transcriptional activity. We first show that the vast majority of reads overlapping LINE-1 elements are derived from pervasive transcription and propose a method to address this issue. We validated the LINE-1 transcription landscape in well-established human cell lines and their cell compartments. Finally, we surveyed LINE-1 activity in a variety of healthy somatic tissues. Although somatic retrotransposition has been mainly studied in the human brain, we found surprisingly little transcriptional activity in most brain regions from adults. Instead, we found LINE-1 transcriptional activity in other somatic tissues consistent with an overall trend of LINE-1 activity in cell with higher turnover.

## Results

Recently amplified LINE-1 subfamilies, such as L1Hs, are frequently discarded from traditional transcript quantification assays due to the insufficient mapping specificity of LINE-1 instances. Before addressing the LINE-1 multi-mappability issue, we quantified the number of reads overlapping LINE-1 subfamilies in thousands of RNA sequencing experiments from human cell lines and healthy primary tissues [31,32]. Figure 1A shows the correlation between the average number of reads mapping to LINE-1 subfamilies in healthy tissues and the number of bases in the reference genome annotated as the respective LINE-1 subfamily (Spearman’s rank correlation rho=0.94, p < 2.2e-16). We find that even ancient LINE-1 subfamilies can have thousands of reads from RNA-seq; for example, reads mapped ten times more frequently to ancient LINE-1 subfamilies, such as L1ME1 and L1M5, than some recently active LINE-1 subfamilies. In fact, most of the LINE-1 reads appeared to derive from subfamilies that are thought to be autonomously inactive (genomic fossils) for millions of years. As an explanation for this counterintuitive result, we hypothesized that this “genomic-transcriptomic” correlation might be indicative of pervasive transcription. In this model, the stochastic nature of RNA polymerase II transcription would drive the creation of RNA fragments proportionally to the number of copies of LINE-1 subfamilies in the whole genome.

**Figure 1.**
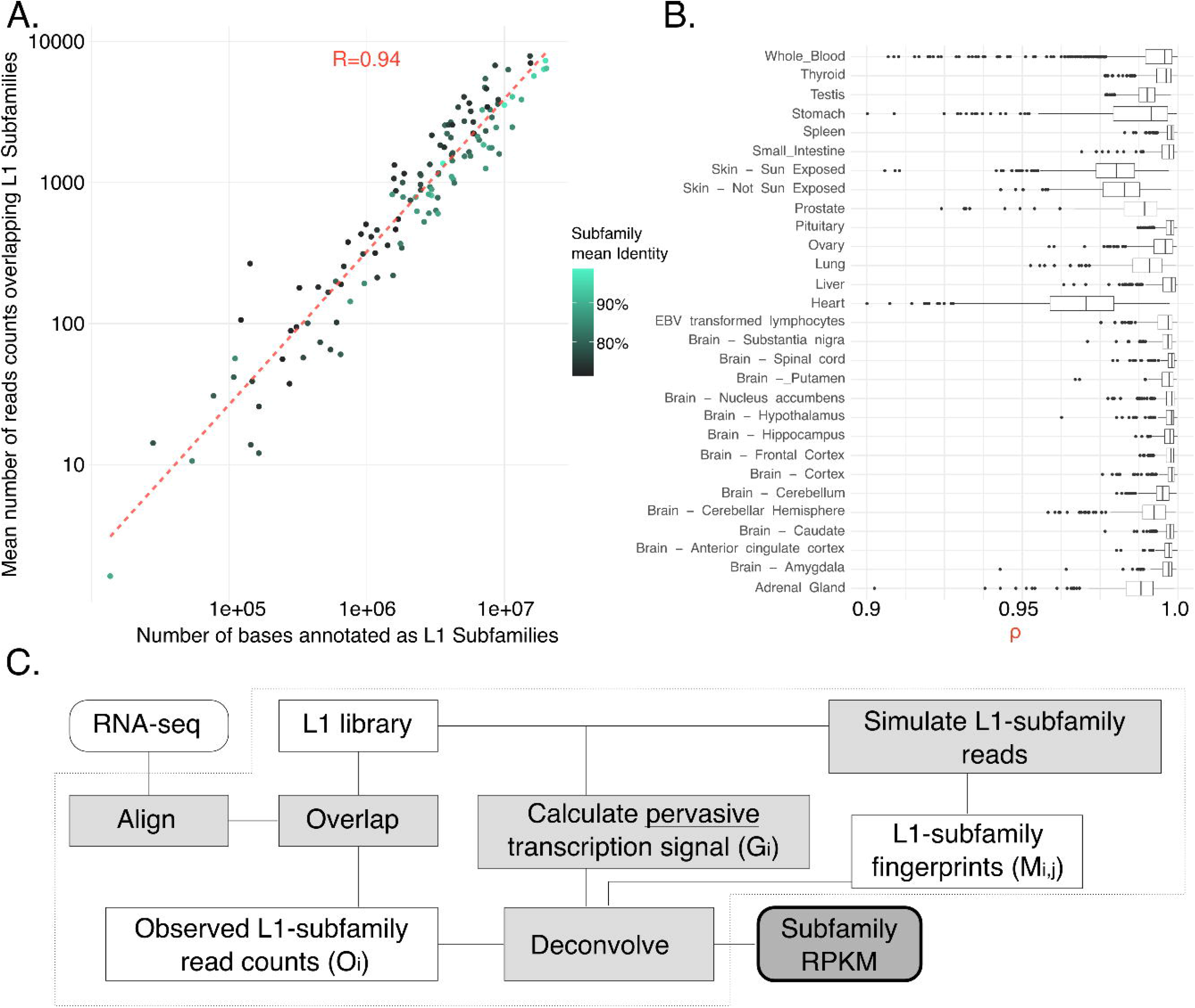
As pervasive transcription is a major factor leading to reads mapping to L1 instances, TeXP functions as an approach to decouple pervasive transcription from autonomous transcription. (A) The number of reads mapped to LINE-1 subfamilies is proportional to the number of bases annotated as the subfamily for most RNA sequencing experiments. Point colors represent the subfamily average identity to LINE-1 consensus. (B) Healthy human tissues show varied distributions of the genomic-transcriptomic correlation. (C) Pipeline chart describes the TeXP approach.

We model pervasive transcription as sufficiently broad transcription component that creates a bias in RNA-seq reads overlapping LINE-1 subfamilies. This bias tends to be correlated to the number of instances (or bases) in the reference genome because pervasive transcription is predicted to create small quantities of reads across large portions of the genome.

As opposed to the broad pervasive transcription model, under the narrow LINE-1 transcription model, we expect that a small number of LINE-1 instances to produce a much higher signal (number of reads) than the lowly expressed instances. Therefore, in order to investigate this issue, we first calculated the number of reads mapping to LINE-1 instances. We observed that, the vast majority of LINE-1 instances have a very small number of uniquely mapped reads (S1A Figure and S1B Figure) suggesting that indeed, pervasive transcription is capable of generating small number of transcripts overlapping many LINE-1 instances across the genome. A typical RNA-seq experiment has approximately 35.000 LINE-1 instance with a single read. In fact, we also observed that, on average, 99.94% of LINE-1 instances with at least one read have very small expression levels (RPKM <= 1).

To test the amount of signal generated by the top expressed instances and the lowly expressed instances we calculated the ratio of reads overlapping the most 10 expressed instances in each RNA-seq experiment and the sum of reads overlapping all instances with less than 10 reads. We find that the summation of the least expressed instances is, on average, one order of magnitude higher than the 10 mostly expressed instances. Together these results suggest that broad pervasive transcription is an important factor when quantifying LINE-1 subfamily transcription level.

We then divided samples by their tissues of origin (Figure 1B) and noticed that some tissues had smaller genomic-transcriptomic correlations, hinting at another confounding signal other than pervasive transcription creating reads overlapping LINE-1 subfamilies. We hypothesized that deviations from a high genomic-transcriptome correlation could be derived from autonomous transcription of the LINE-1 subfamilies (see Methods for details). We then developed a software pipeline, TeXP, that uses mappability signatures from pervasive and simulated LINE-1 subfamilies autonomous transcription to deconvolve reads overlapping LINE-1 elements. TeXP counts the number of reads overlapping recently expanded LINE-1 subfamilies, and calculates the best combination of signatures that explains the observed read counts. Specifically, TeXP regresses the proportion of reads derived from each signal, ensuring sparsity to estimate autonomous transcription of recent LINE-1 subfamilies (L1Hs, L1P1, L1PA2, L1PA3, L1PA4) and remove the effect of pervasive transcription (Figure 1C).

### LINE-1 transcriptional activity in human cell lines

We benchmarked TeXP by estimating the autonomous transcription of LINE-1 subfamilies in RNA sequencing experiments of well-established human cell lines [31]. Figure 2A shows the proportion of reads mapped to LINE-1 subfamilies using a naïve method (left panel) and proportions of reads from each signature using TeXP (right panel). TeXP estimations were also compared to other transposable element quantification pipelines (S2 Figure). In the naïve method (Figure 2A; left panel), cytoplasmic and whole-cell polyadenylated (polyA)+ samples had an enrichment of reads mapping to L1Hs and L1PA2 when compared to whole-cell transcripts without a polyadenylated tail (whole-cell polyA-) and nuclear RNA samples. The enrichment of L1Hs reads was consistent with increased transcription of full-length L1Hs (S3 Figure). The estimates after applying TeXP (Figure 2A; right panel) revealed two major signals in MCF-7 RNA sequencing experiments: pervasive transcription and L1Hs autonomous transcription. The difference between the naïve method and TeXP suggests that reads mapped to ancient LINE-1 subfamilies, such as L1PA3 and L1PA4, are mostly derived from pervasive transcription. TeXP also detected residual L1PA2 transcription in a small number of samples (Figure 2A and S4 Figure). This result is consistent with L1Hs and L1PA2 being the only two LINE-1 subfamilies capable of autonomous transcription and autonomous mobilization in the human germline and tumors [11,33].

**Figure 2.**
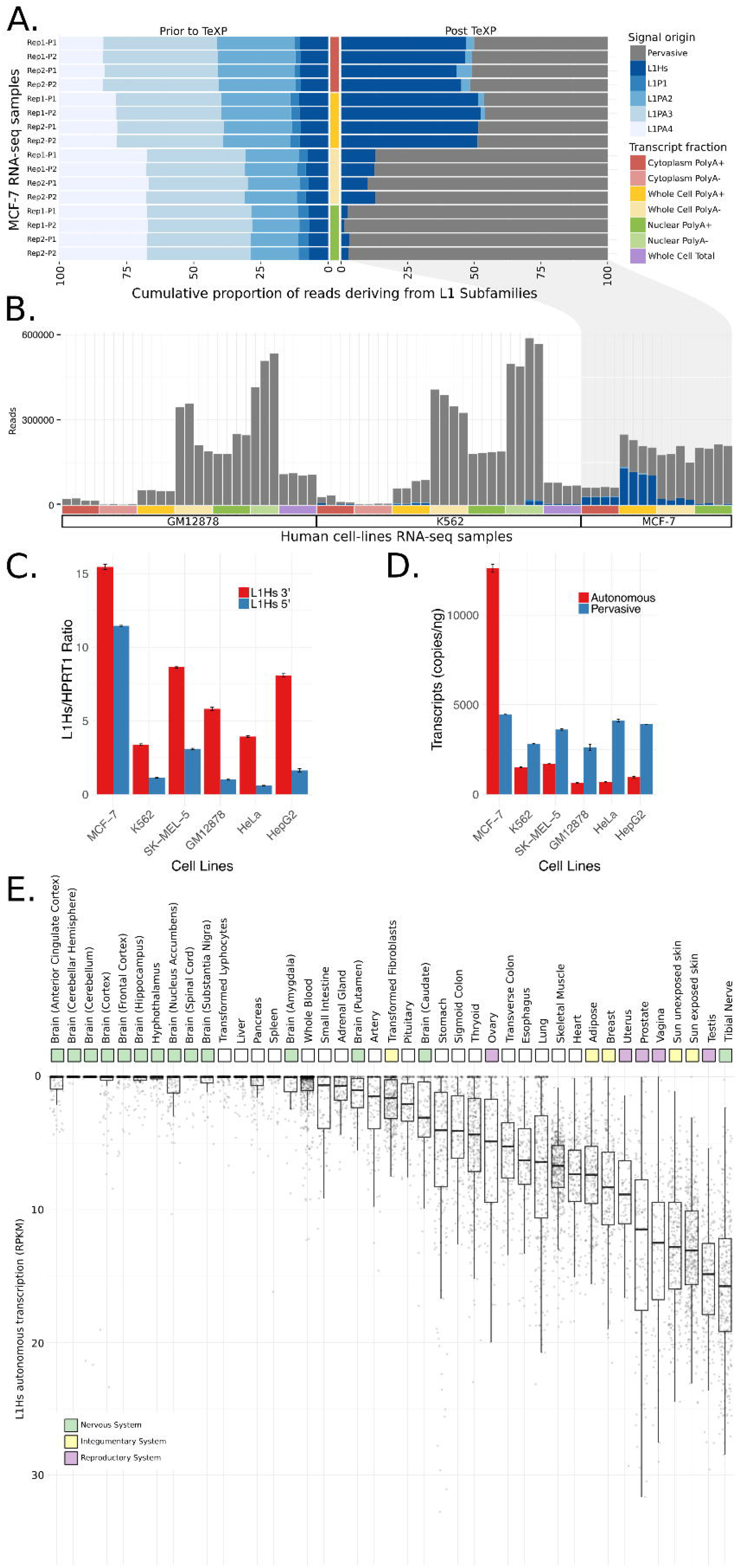
Quantification and validation of L1Hs autonomous transcription in human cell lines. (A) The proportion of reads emanating from pervasive transcription and L1P1, L1PA2, L1PA3, L1PA4, and L1Hs subfamilies in MCF-7 RNA sequencing experiments are shown from the different cell compartments and transcript fractions prior to (left) and after (right) TeXP processing. (B) The absolute number of reads emanating from pervasive transcription and LINE-1 subfamilies are shown across the distinct cell and transcript fractions of the human-derived cell lines GM12878, K-562, and MCF7. (C-D) The quantification of autonomous and pervasive transcripts of L1Hs in the cell lines is shown using ddPCR. (C) The ratio of L1Hs 5’ and 3’ transcripts shows the enrichment of the 3’ end of L1Hs for all cell lines. (D) The absolute quantification of autonomous and pervasive transcripts reveals higher expression of pervasive compared to autonomous transcripts in all cell lines except MCF-7. All data were run in duplicate. All errors bars are mean ± SEM. These data represent two independent experiments. (E) L1Hs autonomous transcription landscape of human healthy primary tissues. Each point is a RNA sequencing experiment, separated by tissue of origin.

MCF-7, a cell line derived from breast cancer, was previously described as having remarkably high levels of L1Hs autonomous transcription [17,24]. The transcriptome of MCF-7 and many other cell lines were carefully and consistently sequenced through the Encyclopedia of DNA elements (ENCODE) project. Leveraging these ENCODE cell line datasets, we assessed L1Hs autonomous transcription in distinct cell compartments (S5 Figure and S6 Figure) [31]. First, we found that MCF-7 whole-cell polyA+ samples had extremely high levels of L1Hs transcription (180.7 RPKM), in agreement with the literature. Selecting whole-cell polyA-samples reduced the signal of L1Hs autonomous transcription by 73% (Figure 2A), suggesting that most of the signal was derived from mature polyA+ LINE-1 transcripts. Furthermore, we tested whether L1Hs transcripts are derived from cytoplasmic (mature) or nuclear (pre-mRNA) portions of the cell. We found that nuclear transcripts were highly enriched for pervasive transcription (autonomous/pervasive ratio 0.02), whereas cytoplasmic transcripts had an autonomous/pervasive ratio similar to transcripts derived from whole-cell polyA+ samples (0.45 and 0.51, respectively – Figure 2A). Together, these results suggest that most of the LINE-1 autonomous transcription signal is derived from mature transcripts in the cytoplasm and only a small fraction of signal is derived from fragmented LINE-1 transcripts in the nucleus. Analyzing other lymphoblastic and cancer-derived cell lines such as GM12878, SK-MEL-5 and K-562 yielded no evidence of L1Hs autonomous transcription in most cell compartments or RNA fractions, despite low levels of L1Hs autonomous transcription in whole-cell polyA+ samples (0, 8.8 and 8.4 RPKM, respectively. Figure 2B and S1 Table).

### Validation of LINE-1 autonomous transcription

To validate the quantification of L1Hs autonomous transcription, we performed droplet digital PCR (ddPCR) to estimate autonomous and pervasive transcription levels on a reference panel of six cell lines: MCF-7, K-562, HeLa, HepG2, SK-MEL-5, and GM12878. For these experiments, we assumed that expression on the 5’ end of the L1Hs transcript can be used as an approximation to autonomous transcription due to the large imbalance of 5’ truncated and full-length copies. The expression on the 3’ end, on the other hand, is an approximation tothe combination of autonomous and pervasive transcription. We initially designed and tested multiple assays targeting different regions of the L1Hs locus and proceeded with the two best performing assays (S2 Table). The first assay targeted ORF1, directly adjacent to the 5’UTR, representing the 5’ end of the transcript. The second assay targeted ORF2 about 1.5 kb upstream of the 3’ UTR, representing the 3’ end of the transcript. We completed the same design process for ORF2 to find the copy numbers of the truncated L1Hs transcripts (i.e., the transcripts missing the 5’ end of L1Hs) (Figure 2C, S3 Table). Since autonomous transcription results in an enrichment full-length transcript of L1Hs, we estimated an approximation to the level of pervasive transcription by subtracting expression of the 5’ end (ORF1) from the 3’ end (ORF2).

Figure 2D shows the relative quantification of L1Hs transcripts in these four cell lines using the *HPRT1* 5’ end as a reference. The ddPCR analysis detected 12,600 copies of full-length transcripts/ng in MCF-7 cells. In agreement with our *in-silico* result, K562 and SK-MEL-5 had 1,512 and 1,708 copies of full-length transcript/ng, respectively. For the GM12878 cell line, we expected to find no autonomous expression of L1Hs; however, our ddPCR assays detected low levels of autonomous transcription of L1Hs (655 copies of full-length transcript/ng; Figure 2D, S3 Table). Overall, the quantification of L1Hs autonomous transcription using ddPCR was highly correlated with the quantification using TeXP (Spearman correlation, rho=0.99, p-value=3.803e-06, S7 Figure). This suggests that TeXP can remove most of the noise derived from pervasive transcription, although it is insensitive to samples with little LINE-1 autonomous transcription (S8 Figure). To address TeXP sensitivity in relation to noise we first tested TeXP under an ideal experimental setup. In this simulation, the number of observed reads overlapping L1 subfamilies is a simple combination of known proportions of these signals (i.e. pervasive and autonomous transcription). For example, we simulate read counts where 30% of the reads derive from LIHs autonomous transcription and the remaining 70% derive from pervasive transcription. We use simulated read counts as input to TeXP and calculate the root mean square error (rmse) between the known proportions and estimated proportions. As observed in S15 Figure (solid line), under this condition, the TeXP estimations of read counts is nearly identical to the simulated read counts (median(rmse) = 0.0003). We also tested the effect of noise in the TeXP estimations. To that end, we modeled the noise component as a Poisson process. For these simulations, the Poisson noise was added to read counts deriving from of known proportions of signals as described above. We tested two scenarios, one with noise equivalent to 10% of all observed reads (dashed line) and 20% of the of all observed reads (long dashed lines). In these conditions, the median TeXP RMSEs were respectively 0.014 and 0.028. Overall our observations suggest that, under low noise conditions, TeXP should be able to detect autonomous L1Hs transcription even when L1Hs has low transcription levels. However, we also describe the TeXP capacity to detect low levels of L1Hs autonomous transcription degrades with higher levels of noise in RNA-se data. In extreme situations, such as when 20% of the read counts are derived for a Poisson process, the RMSE can be up to 0.083.

### Landscape of LINE-1 subfamily transcription in healthy primary tissue and cells lines

Researchers have long thought that LINE-1 instances are completely silenced in most healthy somatic cells. LINE-1 is silenced by the methylation of its promoter [19], which should preclude the transcription of mature LINE-1 mRNAs in healthy somatic tissue. To test whether LINE-1 subfamilies are completely silenced in somatic tissue, we analyzed LINE-1 transcription in 7,429 primary tissue samples from the Genotype-Tissue Expression (GTEx) project [32] (S4 Table). Similar to the cell lines, we found that L1Hs was autonomously transcribed; L1P1, L1PA2, L1AP3, and L1PA4 only had residual or spurious autonomous transcription in healthy tissues (S9 Figure). Furthermore, we found that pervasive transcription was the major signal in most RNA sequencing datasets, accounting for 91.7%, on average, of the reads overlapping LINE-1 instances (S10 Figure and S14 Figure). Overall, healthy tissues had a narrower range of L1Hs autonomous transcription levels than cell lines, with the peak transcription level of 47 RPKM (Figure 2E) versus 180 RPKM in the cell lines (S1 Table). We found no or very little (<1 RPKM) evidence of L1Hs autonomous transcription in 2,520 (34.3%) of the GTEx RNA sequencing experiments from primary tissues. Together, these results indicate that L1Hs is broadly transcribed in some healthy somatic tissues. Therefore, if post-transcriptional regulatory constraints do not completely silence LINE-1 activity, one could expect that LINE-1 to play an important role in creating genetic diversity across somatic cells within an individual.

We then compared the landscape of LINE-1 subfamily transcription in Epstein-Barr virus (EBV) immortalized cell lines and their corresponding primary tissue to understand the changes induced by cell line immortalization. EBV immortalization causes drastic changes in the expression of cell cycle, apoptosis, and alternative splicing pathways [34–36]. Overall, we found that EBV-transformed cell lines derived from different tissues (lymphoblastic and fibroblastic) had distinct patterns of L1Hs autonomous transcription; lymphoblast (blood-derived) cell lines had no or little autonomous transcription of L1Hs (S11 Figure) with approximately 84% of samples having an estimated RPKM equal to zero, whereas fibroblastic (skin-derived) cell lines consistently had higher levels of L1Hs autonomous transcription (median 1.5 RPKM) with 58.7% of samples having an RPKM higher than 1. In general, EBV-immortalized cell lines reflected their tissue of origin. While most (74.6%) of the whole blood samples had no transcriptional activity of L1Hs, only one sample from skin had an L1Hs autonomous transcription level below 1 RPKM. We further selected patients with both primary and EBV-transformed cell lines to assess whether the EBV transformation could change L1Hs autonomous transcription. We found that both skin cells and lymphocytes had a drastic down-regulation of L1Hs autonomous transcription (S12 Figure). This finding suggests that EBV-transformed cell lines partially preserve the L1Hs transcription level from their tissue of origin, potentially explaining why fibroblast-derived induced pluripotent stem cells support higher levels of LINE-1 retrotransposition [37].

Human solid tumors display increased levels of LINE-1 activity [38–40]. In order to assess if this increase is a result from pervasive transcription or an increase in autonomous transcription in LINE-1 we run TeXP in RNA-seq from healthy thyroid samples and solid thyroid tumors. We than calculate the distribution of number of reads deriving from pervasive and autonomous transcription. We first observed that there indeed a significant difference in the number of reads from LINE-1 elements (S13 Figure). We also found that compared to healthy thyroid samples, tumor samples display higher levels of autonomous transcription and lower levels of pervasive transcription suggesting that autonomous transcription LINE-1 is driving most of the increase in LINE-1 expression in thyroid tumor samples.

Human tissues show remarkable variability of L1Hs autonomous transcription. We found that L1Hs autonomous transcription is inversely correlated to the time it takes cells to divide (cell turnover rate; Pearson correlation: cor=-0.6668968; p-value=0.04). No correlation was found between cell turnover and pervasive transcription (Pearson correlation: cor=0.3983474; p-value = 0.2883). Tissues suggested to have low cell turnover, such as the human brain [41], are amongst the tissues with the lowest levels of L1Hs autonomous transcription (Figure 2E). In particular, the human cerebellum, which has no transcription of L1Hs, is likely to have strong repression of L1Hs autonomous transcription. This result seems to contradict the literature that suggests that the human brain supports high levels of somatic LINE-1 retrotransposition; however, most of these studies were based on neuro precursors that correspond to the early development stage of the human brain [15,42–44]. Conversely, brain samples extracted from the striatum, putamen, and caudate, all regions associated with the basal ganglia, had higher levels of L1Hs autonomous transcription compared to other brain regions (T-test basal ganglia vs. all other brain tissues, t = −7.0943; p value = 9.867e-12 – Figure 2E); importantly, these levels were still low compared to other tissues. Other tissues with low cell turnover rates, such as liver, pancreas, and spleen, also showed very little or no autonomous transcription of L1Hs (91.2%, 82.9%, 88.9% of samples, respectively, had a L1Hs RPKM < 1 – Figure 2E). Conversely, germinative tissues have been proposed to support somatic activity of L1Hs elements [45]. Our results suggest that this trend is more general, and most tissues associated with the reproductive system sustain higher levels of L1Hs autonomous transcription (Figure 2E). In addition, we found that the tissues with the highest levels of L1Hs autonomous transcription were enriched for high cell turnover; these included the nerve (tibia), skin (both exposed and not exposed to the sun), prostate, lung, and vagina (Figure 2E).

## Discussion

Prior to this study, the effects of pervasive transcription on the estimates of transposable elements activity were largely ignored. Here, we showed that most of the RNA-seq reads matching to LINE-1 instances derive from pervasive transcription, highlighting the importance of these effects. In order to account for the effect of pervasive transcription on the quantification of LINE-1 activity we developed TeXP, a method that uses the widespread nature and the mappability signatures of LINE-1 subfamilies to account for and remove the effects of pervasive transcription. We compared TeXP estimates to other strategies such as naïve counts and others established methods such as SalmonTE [46] and TETranscript [47] (S2 Figure). Our estimations suggest that the pervasive transcription component is frequently missed and can be a confounding factor in some quantifications, for example, we observed that over 75% of reads mapping to ancient LINE-1 subfamilies derive from inactive subfamilies (Figure 2A and S14 Figure).

We used TeXP to perform a comprehensive analysis of LINE-1 transcriptional activity across different cell types and somatic tissues. Previous studies suggest that LINE-1 is active in germline and tumor cells, but not in most normal somatic cells with the exception of hints of activity in neuro-precursor cells [21]. Somatic mosaicism of transposable elements, in particular LINE-1, has been carefully characterized in the human brain and, despite some disagreement on the exact rate that LINE-1 retrotranspose in the human brain, it is clear that LINE-1 frequently create somatic copies in the human brain, in particular, in neuroprecursor, cortex and caudate nucleus cells [15,48,49]. The somatic mobilization in other tissues, nonetheless, remain obscure and lack a systematic investigation. A first step is to comprehensive characterize other tissues in terms of their LINE-1 transcriptome activity. Surprisingly, we found that LINE-1 was active in many healthy human tissues, particularly in epithelial cells. As we only detected a limited amount of LINE-1 activity in adult brain cells, our findings are in agreement with LINE-1 activity correlating with cell proliferation rate.

We validated many aspects of TeXP using ddPCR probes designed to quantify pervasive and autonomous transcription of L1Hs across human cell lines. These results show that our method lacks the sensitivity to measure autonomous transcription. In particular, TeXP underestimates L1Hs transcription levels when the signal-to-noise ratio is small. One could imagine using unique regions of LINE-1 instances after removing the pervasive transcription signal to improve the transcriptional quantification. However, removing pervasive transcription from individual instances is not trivial and should be carefully investigated.

As a tool, TeXP could be useful in several scopes beyond this study. Pervasive transcription should also affect the quantification of other transposable elements and repetitive regions accounting for large portions of the human genome. Our method could be further used to estimate the autonomous transcription levels of pseudogenes, SINEs (ALUs) and HERVs. Furthermore, TeXP could be used in model organisms to distinguish the effects of pervasive transcription beyond humans. The mouse genome, for example, has evidence of higher rates of retrotransposition but little is known about the activity of LINE-1 in somatic cells. Moreover, some of the results we describe here could be extended and uncover important biological insights. For example, assays such as induced pluripotent stem cells clones [50], RC-seq [51] and L1-seq [52], among others, could be used to carefully characterize of the rate of somatic retrotransposition in tissues with higher rates of autonomous transcription of L1Hs. Additionally, TeXP could be used to further investigate the biases found in individuals from different ancestral backgrounds [53]. The TeXP approach to quantify transcriptional activity by removing pervasive transcription could also be expanded to investigate the activity of LINE-1 during embryonic development or in pathological tissues, such as tumors.

## Materials and Methods

### Modeling pervasive and autonomous LINE-1 transcription

TeXP models the number of reads overlapping L1 elements as the composition of signals deriving from pervasive transcription and L1 autonomous transcripts from distinct L1 subfamilies.

Our model proposes that the number of reads overlapping L1Hs instances as described by the **Equation 1**:

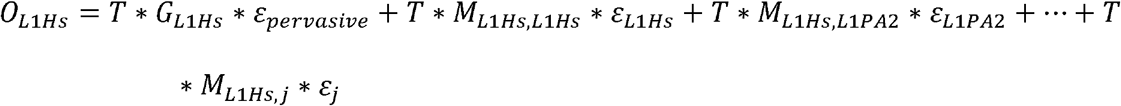

Where *O*_*L*1*Hs*_ is the observed number of reads mapping to L1Hs, T is the total number of reads mapped to L1 instances, *G*_*L*1*Hs*_ defines the proportion of L1 bases in the genome annotated as L1Hs, *ε_pervasive_* is the percentage of reads emanating from pervasive transcription, M is the mappability fingerprint (defined bellow) that describes what is the proportion of reads emanating from the signal *j* ∈ {*L*1*Hs*,*L*1*P*1,*L*1*PA*2,*L*1*PA*3,*L*1*PA*4} that maps to L1 subfamily *i* ∈ {*L*1*Hs*,*L*1*P*1,*L*1*PA*2,*L*1*PA*3,*L*1*PA*4} and *ε* is the percentage of reads emanating from the autonomous transcription L1 Subfamily *j*. This model can be further generalized as the **Equation 2**:

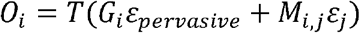

We selected these five LINE-1 subfamilies based on the rates of cross-mappability of simulated data (S3 Figure). In particular, we are interested in removing the effect of pervasive transcription from the estimates of L1Hs autonomous transcription. We simulated reads from L1Hs transcripts we observed that most (>90%) of the reads emanating from L1Hs autonomous transcription map to the L1Hs, L1PA2, L1PA3, L1PA4 and L1P1 subfamilies. Older subfamilies such as L1PA5 and L1PA6, for example, correspond to less than approximately 5% of L1Hs cross-mappable reads.

In contrast to L1Hs which only a fourth of the reads map back to L1Hs, older elements such as L1PA5 and L1PA6 have a much higher self-mapping rates, 60% and 70% respectively. Therefore, these subfamilies should be less affected by confounding factors deriving from pervasive transcription.

The number of reads mapped to each subfamily *O_i_* is measured by analyzing paired-end or single-end RNA sequencing experiments independently. TeXP extracts basic information from fastq raw files such as read length and quality encoding. Fastq files are filtered to remove homopolymer reads and low quality reads using in-house scripts and FASTX suite (http://hannonlab.cshl.edu/fastx_toolkit/). Reads are mapped to the reference genome (hg38) using bowtie2 (parameters: --sensitive-local -N1 --no-unal). Multiple mapping reads are assigned to one of the best alignments. Reads overlapping LINE-1 elements from Repeat Masker annotation of hg38 are extracted and counted per subfamily. The total number of reads T is defined as 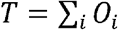.

### Pervasive transcription and mappability fingerprints of L1 subfamily transcripts

Pervasive transcription is defined as the transcription of regions well beyond the boundaries of known genes [27]. We rationalized that the signal emanating from pervasive transcription would correlate to the number of bases annotated as each subfamily in the reference genome (hg38). We used Repeat Masker to count the number of instances and number of bases in hg38 annotated as the subfamily *i* ∈ {*L*1*Hs*,*L*1*PA*2,*L*1*PA*3,*L*1*PA*4,*L*1*P*1}. We define *P_i_* as the proportion of LINE-1 bases annotated as the subfamily *i* in the **Equation 3**:

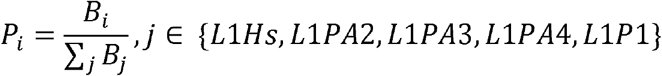

On the other had mappability fingerprints, which represents how reads deriving from LINE-1 transcripts would be mapped to the genome, are created by aligning simulated reads deriving from putative L1 transcripts from each L1 subfamily. For each L1 subfamily, we extract the sequences of instances based on RepeatMasker annotation and the reference genome (hg38). Read from putative transcripts are generated using wgsim (https://github.com/lh3/wgsim - parameters: −1 [RNA-seq mean read length] –N 100000 -d0 −r0.1 -e 0). One hundred simulations are performed and reads are aligned to the human reference genome (hg38) using the same parameters described in the model session. The three-dimensional count matrix *C* is defined as the number of reads mapped to the subfamily *i* ∈ {*L*1*Hs*,*L*1*PA*2,*L*1*PA*3,*L*1*PA*4,*L*1*P*1} emanating from the set of full-length transcripts *j* ∈ {*L*1*Hs*,*L*1*PA*2,*L*1*PA*3,*L*1*PA*4,*L*1*P*1} in the simulation *k*. The matrix M is defined as the median percentage of counts across all simulations as in **Equation 4**:

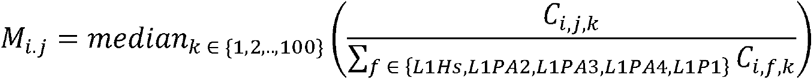

We tested whether different aligners yield different mappability fingerprints. BWA, STAR, and bowtie2 yielded similar results (S15 Figure). As L1 transcripts are not spliced, we decided to integrate bowtie2 as the main TeXP aligner. We further tested the effect of read length on L1Hs subfamily mappability fingerprints (S16 Figure). To counter the effects of distinct read lengths TeXP constructs L1 mappability fingerprints libraries based on fastq read length.

We simulated reads emanating from their respective L1 subfamily transcripts and aligned these reads to the human reference genome creating a mappability fingerprint for each L1 subfamily (S3 Figure). When we analyzed the L1 subfamily mappability fingerprints we observed that younger L1 subfamilies tend to have more reads mapped to other L1 subfamilies. For example, we find that only approximately 25% of reads from L1Hs (the most recent – and supposedly active L1) maps back to loci annotated as L1Hs. While older subfamilies such as L1PA4, have a higher proportion of reads mapping back to its instances (~70% - S3 Figure).

### The hidden variables *ε* and *ϵ*

The known variables *O_i_*, T, the vector *P_i_* the mappability fingerprint matrix *M_i.j_* are used to estimate the signal proportion *ε* and *ϵ* in **Equation 2** by solving a linear regression. We used lasso regression (L1 regression) to maintain sparsity. We used the R package penalized ([54] - parameters: unpenalized=~0, lambda2=0, positive=TRUE, standardize=TRUE, plot=FALSE, minsteps=10000, maxiter=1000).

### TeXP availability

TeXP was developed as a combination of bash, R and python scripts. The source code is available at https://github.com/gersteinlab/texp. A docker image is also available for users at dockerhub under fnavarro/texp.

### GTEx raw RNA sequencing data

Raw RNA sequencing datasets from healthy tissues were obtained from Database of Genotypes and Phenotypes (DB-Gap - https://dbgap.ncbi.nlm.nih.gov) accession number phs000424.v6.p1.

### ENCODE raw RNA sequencing data

Raw RNA sequencing data from cell lines were obtained from the ENCODE data portal (https://www.encodeproject.org/search). We selected RNA-seq experiments from immortalized cell lines with multiple cellular fractions and transcripts selection experiments. Accessions and cell lines are available in TableS1.

### Cell turnover rate

Mutation load and cell turn-over rate were extracted from the compilation of somatic mutation rate in Tomasetti et al [55].

### Pervasive versus Autonomous transcription of L1Hs transcripts

More ancient elements such as DNA transposons and LINE-2 have been shown to be primarily transcribed pervasively, hitchhiking the transcription of nearby autonomously transcribed regions [32]. Therefore, we tested whether our estimation of L1Hs transcription level correlated with genes containing or adjacent to L1 Hs instances. We found no significant difference between the correlation distribution of a random set of genes and genes with L1Hs in exons or introns or within 3kb upstream or 3kb downstream of L1Hs. This finding indicates that our estimation of L1Hs autonomous transcription is not significantly influenced by non-autonomous L1Hs transcription adjacent or contained by protein-coding genes’ loci. Furthermore, we tested if and enrichment of pervasive transcription deriving from intronic regions would create a background signal distinct from the pervasive transcription derived from a whole genome model. We correlated the number of LINE-1 instances from each subfamily in intergenic and intronic regions based on GENCODE v29 and found a statistically significant correlation between the number of instances in both regions (Spearman corr=0.979057, p-value < 2.2e-16 - S17 Figure).

### Cell Culture and Culture Conditions

All the cell lines used in this study were obtained from the American Type Culture Collection (ATCC) (Manassas, VA, USA). MCF-7 cells were cultured in Dulbecco’s Modified Eagle Medium: Nutrient Mixture F-12 (DMEM/F12; Gibco). HeLa, SK-MEL-5, and HepG2 cells were cultured in Dulbecco’s Modified Eagle Medium (DMEM; Gibco). K562 and GM12878 cells were cultured in RPMI 1640 (Gibco). All cell culture media were supplemented with 10% fetal bovine serum (FBS) (Atlanta Biologics) and 1% penicillin/streptomycin (Fisher Scientific). All cells were cultured and expanded using the standard methods.

### RNA Extraction and cDNA Synthesis

RNA was extracted using the RNeasy PLUS Mini Kit and the QIAshredders (Qiagen) following the manufacturer’s protocol. All samples were treated with DNase I (New England BioLabs Inc.) to remove any remaining genomic DNA. RNA concentration was determined by Qubit 2.0 Fluorometer (Invitrogen). RNA quality was determined by Nanodrop (Thermo Scientific) and 2100 BioAnalyzer with the Agilent RNA 6000 Nano kit (Agilent Technologies). Approximately 5 μg of RNA was used for synthesis of the cDNA using the iScript Advanced cDNA Synthesis Kit (Bio-Rad). The final cDNA product was quantified and a working solution of 10 ng/μL was prepared for the subsequent studies.

### Droplet Digital PCR (ddPCR)

Droplet Digital PCR (ddPCR) System (Bio-Rad Laboratories) was utilized to quantify the L1Hs transcript expression in the cell lines described above. Since L1Hs is a highly repetitive and heterogeneous target, we had initially designed and tested a panel of primers and probes that targeted the 5’ untranslated region (5’UTR), the open reading frame 1 (ORF1), the open reading frame 2 (ORF2), and the 3’ untranslated region (3’UTR) of the L1Hs locus, respectively. After a pilot screening study, we selected the two assays covering ORF1 and ORF2, which not only exhibited overall better performance, but also could help us to distinguish autonomous and pervasive L1Hs transcriptions. We also designed two reference assays on the housekeeping gene *HPRT1*, which targeted the 5’ and 3’ ends of the transcript, respectively (S2 Table). All the ddPCR primers and probes were designed based on the human genome reference hg19 (GRCh37) and synthesized by IDT (Integrated DNA Technologies, Inc. Coralville, Iowa, USA).

The ddPCR reactions were performed according to the protocol provided by the manufacturer. Briefly, 10ng DNA template was mixed with the PCR Mastermix, primers, and probes to a final volume of 20 μL, followed by mixing with 60 μL of droplet generation oil to generate the droplet by the Bio-Rad QX200 Droplet Generator. After the droplets were generated, they were transferred into a 96-well PCR plate and then heat-sealed with a foil seal. PCR amplification was performed using a C1000 Touch thermal cycler and once completed, the 96-well PCR plate was loaded on the QX200 Droplet Reader. All ddPCR assays performed in this study included two normal human controls (NA12878 and NA10851) and two mouse controls (NSG and XFED/X3T3) as well as a no-template control (NTC, no DNA template). All samples and controls were run in duplicates. Data was analyzed utilizing the QuantaSoft™ analysis software provided by the manufacturer (Bio-Rad). Data were presented in copies of transcript/μL format which was mathematically normalized to copies of transcript/ng to allow for comparison between cell lines.

### Reference house-keeping gene (HPRT1)

We designed two assays targeting the 5’ and 3’ ends of the *HPRT1* transcript, respectively, and used as the reference controls in this study (S3 Table). The reference gene expression level was found to be constant within each cell line, but varied between cell lines. In addition, while 4 of the 6 cell lines had similar 5’ and 3’ end expression, K562 and GM12878 both had increased 3’ end expression. This could be from different isoforms being expressed with different frequencies^3^. For the 5’ end expression of *HPRT*, SK-MEL-5, GM12878, and HepG2 were all around 600 copies of transcript/ng. The remaining were all around 1200 copies of transcript/ng. When looking at the 3’ end expression, we found that SK-MEL-5 and HepG2 were around 750 copies of transcript/ng, while MCF-7, GM12878, and HeLa were around 1350 copies of transcript/ng, and K562 was close to 1800 copies of transcript/ng. The slight difference between the 5’ end and the 3’ end expression levels in the same cell line could be explained by a potential 3’ end bias in the cDNA synthesis. However, all the reference assays were consistent between experiments and did not affect the target expression.

## Acknowledgements

This work was supported in part by a grant from the National Human Genome Research Institute (grants U24HG009446 and U41HG007497) and the National Cancer Institute of the NIH (grant P30CA034196). C.L. is a distinguished Ewha Womans University Professor, supported in part by the Ewha Womans University Research grant of 2016.

